# Evidence that alpha blocking is due to increases in system-level oscillatory damping not neuronal population desynchronisation

**DOI:** 10.1101/729723

**Authors:** David T. J. Liley, Suresh D. Muthukumarswamy

**Affiliations:** Department of Medicine, The University of Melbourne, Parkville VIC 3010, Australia; Centre for Human Psychopharmacology, Swinburne University of Technology, Hawthorn VIC 3122, Australia; Department of Pharmacy and Pharmacology, The University of Auckland, Auckland, New Zealand

**Keywords:** electroencephalogram, magnetoencephalogram, alpha rhythm, alpha blocking, alpha desynchronisation

## Abstract

The attenuation of the alpha rhythm following eyes-opening (alpha blocking) is among the most robust features of the human electroencephalogram with the prevailing view being that it is caused by changes in neuronal population synchrony. To further study the basis for this phenomenon we use theoretically motivated fixed-order Auto-Regressive Moving-Average (ARMA) time series modelling to study the oscillatory dynamics of spontaneous alpha-band electroencephalographic activity in eyes-open and eyes-closed conditions and its modulation by the NMDA antagonist ketamine. We find that the reduction in alpha-band power between eyes-closed and eyes-open states is explicable in terms of an increase in the damping of stochastically perturbed alpha-band relaxation oscillatory activity. These changes in damping are putatively modified by the antagonism of NMDA-mediated glutamatergic neurotransmission but are not directly driven by changes in input to cortex nor by reductions in the phase synchronisation of populations of near identical oscillators. These results not only provide a direct challenge to the dominant view of the role that thalamus and neuronal population de-/synchronisation have in the genesis and modulation of alpha electro-/magnetoencephalographic activity but also suggest potentially important physiological determinants underlying its dynamical control and regulation.

## 1. Introduction

The resting alpha rhythm (8–13 Hz) is arguably the most ubiquitous cortical electroencephalographic rhythm. Spontaneous alpha-band activity can be recorded from all areas of intact cortex and in many cases shows a remarkable attenuation in response to a broad range of stimuli and behaviours (Chang et al., 2010). Originally referred to as the Berger effect (Adrian and Matthews, 1934), in honour of its discoverer Hans Berger (Berger, 1929), it is now more commonly referred to as alpha blocking or desynchronisation. Classically clear alpha-band oscillations can be seen in occipital cortex (the classical alpha rhythm), central Rolandic areas (the so called mu rhythm) and mid-temporal regions (the so called ‘third rhythm’)(Chang et al., 2010). However, despite many decades of investigation involving the alpha rhythm and its physiological basis and cognitive correlates, little certainty has been achieved regarding its generative mechanisms and functional roles (Lozano-Soldevilla, 2018).

The received view regarding the physiological mechanisms responsible for alpha involves a central role of the thalamus. The earliest theories suggested that it was activity in the thalamus that was ‘pacing’ or driving overlying cortical tissue through excitatory thalamo-cortical afferents (Andersen and Andersson, 1968). Such a view has been refined in the last 20 years by positing that it is through a complex interaction of intrinsic high-threshold bursting and gap junctions by which thalamic, and thereby cortical, alpha is generated (Hughes and Crunelli, 2005). Concordant with such refinements have been the development of theories that account for alpha in terms of emergent reverberant feedback activity between thalamus and cortex (Vijayan et al., 2013; Jones et al., 2009; Robinson et al., 2002). These thalamic-based theories explain the phenomena of alpha blocking in terms of altered thalamic (driving) activity based upon empirical reports indicating that during occipital alpha rhythm episodes there is an increased activation of thalamic areas as measured by blood oxygenation or glucose metabolism (Feige et al., 2005; Goldman et al., 2002; Sadato et al., 1998). These findings reiterate direct electrocortical recordings in cats and dogs in which alpha rhythms in the visual system and thalamus are reported to occur simultaneously with the corresponding cortical rhythms (Chatila et al., 1993, 1992; Lopes da Silva et al., 1980, 1973).

However, there exist a number of macroscopic mean field theories that suggest that the alpha rhythm might better be explained in terms of the reverberant activity within cortical populations of neurons (Liley et al., 2012; Deco et al., 2008). These models suggest that the alpha rhythm emerges from feed-forward and feed-back axo-synaptic connectivity between excitatory and inhibitory *populations* of neurons. While these models are typically formulated as coupled non-linear stochastic partial differential equations they can be linearised in order to generate physiologically plausible noise-driving spectra. In the case of the well-known non-linear model of Liley et al (Coombes, 2010; Liley et al., 2003, 2002; Steyn-Ross et al., 1999) the fluctuation spectra so obtained typically evince ‘1/*f*’ behaviour at low frequencies and a clear resonance centred at alpha frequencies for a range of physiologically plausible parameterisations. Mechanistically the Liley model suggests that alpha-band oscillatory activity arises principally as a consequence of synaptic feed-forward and feed-back delays between excitatory and inhibitory neuronal populations that arise due to the characteristic time courses of the respective postsynaptic potentials (Hartoyo et al., 2019).

Importantly such an approach suggests that alpha blocking can be accounted for in terms of the parametric damping of stochastic alpha-band oscillatory activity. In particular, it suggests that the well-described reductions observed in alpha power between eyes-closed (EC) and eyes-open (EO) conditions might be understood in terms of an increase in the damping (i.e. the rate at which amplitude decreases with time) of randomly excited alpha-band oscillatory activity. Such a hypothesis implies that the frequency domain alpha resonance will broaden, an effect not compatible with alpha power decreases being driven by reductions in neuronal population oscillatory synchrony and/or cortical driving. This can be easily seen by taking the Fourier Transform of a damped oscillation of the form Θ(*t*)*e^−γt^* cos(2*πft*), where Θ(*t*) is the Heaviside step function, *γ* is damping and *f* is the oscillatory frequency (see Fig. 1 for illustrative purposes).

**Figure 1:**
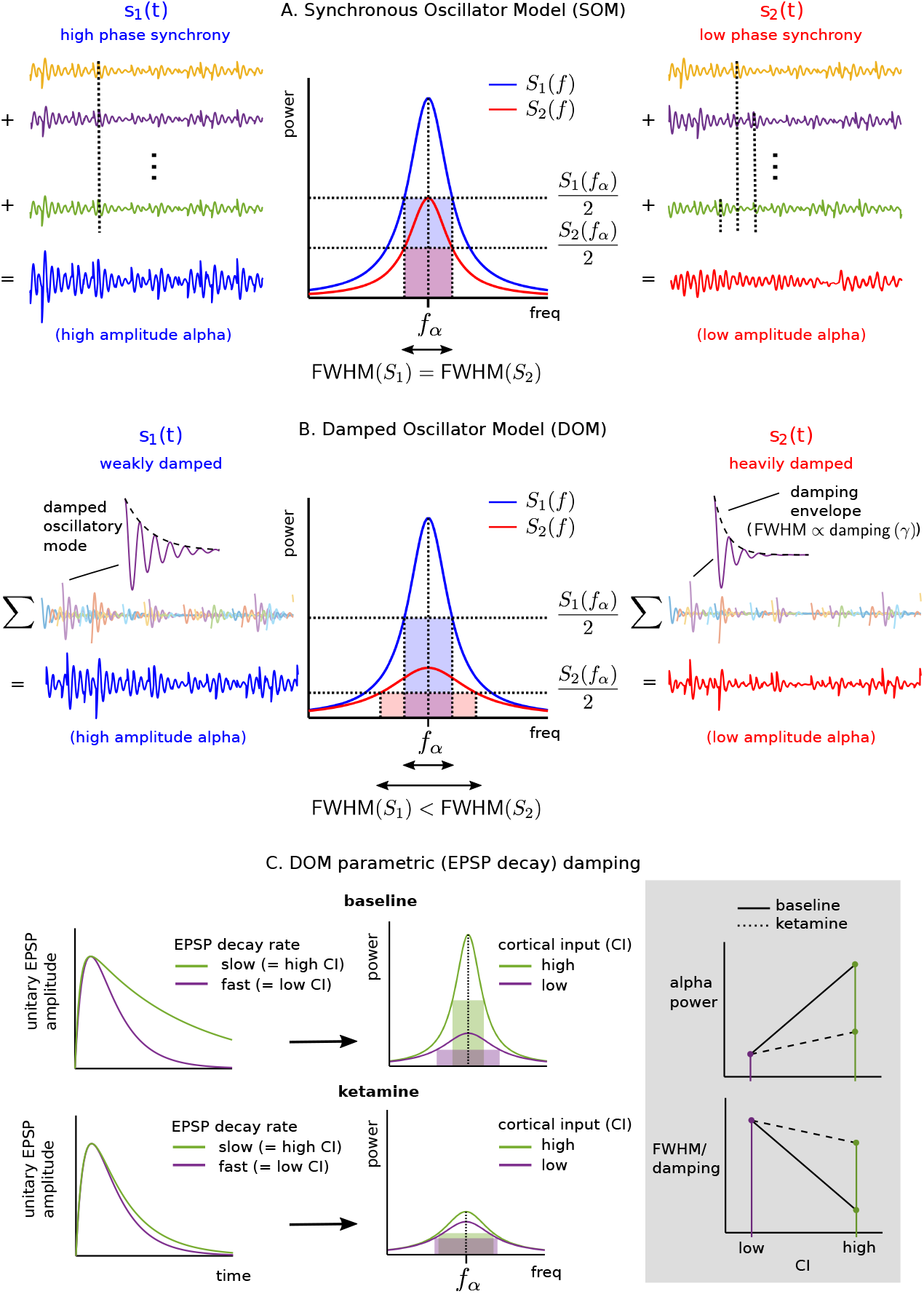
A - Cartoon depiction of the predicted effects that two different models developed to explain variations in the amplitude of alpha (8–13 Hz) EEG activity have on the spectral shape of alpha-band oscillatory activity. A - The Synchronous Oscillator Model (SOM) posits that variations in the phase synchrony of a population of near identical alpha oscillators causes variations in the amplitude of alpha-band activity. Because the shape of a signal’s power spectrum will be invariant to changes in signal phase the full-width half-maximum (FWHM) of the central alpha (*f_α_*) resonance will be invariant to any variations in the amplitude of alpha activity. B - The Damped Oscillator Model (DOM), in contrast, hypothesises that changes in the decay rate (damping, *γ*) of stochastically perturbed alpha oscillatory activity is the source of variations in the amplitude of alpha-band EEG activity. Increases in the decay/damping of such alpha oscillatory modes will broaden the associated spectral resonance and thus increase its FWHM. In this simple model changes in the variance of the stochastic driving process (cortical input, CI) will have no effect on the FWHM. C - The model of Liley et al. (2002) predicts that the NMDA-mediated prolongation of the decay of the unitary excitatory postsynaptic potential will “sharpen” the alpha resonance i.e. reduce the alpha spectral FWHM, by decreasing the damping of stochastically perturbed alpha oscillatory modes. Such NMDA mediated effects are hypothesised to be driven by changes in CI and therefore it is predicted that the addition of the NMDA antagonist ketamine will act to reduce the magnitude of changes in alpha power and FWHM/damping as a function of CI.

From this neurobiologically informed theoretical perspective it is found that a wide range of parameters determine the frequency and damping of alpha-band activity. Of particular interest are those parameters that directly modulate cortical excitation - excitatory thalamo-/cortico-cortical driving and the time course of the feedback excitatory postsynaptic potentials. Slower excitatory postsynaptic potentials (EPSPs) and increases in excitatory thalamo-/cortico-cortical driving (i.e. cortical input) are found to produce more sharply resonant alpha-band activity (Liley et al., 2002). While the predicted effects of changes in cortical input accord well with previous empirical neurobiological explanations for modulations of alpha power the neurobiological basis for changes in EPSP time course appear much less obvious, that is until we consider the properties of the voltage-dependent ionotropic NMDA receptor channel (Hestrin et al., 1990). Because excitatory synapses are composed of both ‘fast’ AMPA/kainate and ‘slow’ NMDA ionotropic receptors the net effect of postsynaptic membrane depolarisation on the unitary EPSP is to essentially prolong its decay phase as NMDA mediated ionic currents become activated (see Fig. 1C). Therefore, cortical input is predicted to not only directly drive changes in the damping of population alpha activity but it is also predicted to indirectly drive changes in alpha damping by modulating changes in the EPSP time course through activity-dependent NMDA-mediated mechanisms.

Therefore, on the basis of the forgoing theoretical arguments, our aim is to empirically evaluate the following hypotheses:

1. Alpha-blocking will be associated with changes in the damping of alpha oscillatory modes (also referred to as alpha poles). In particular, EO alpha will be more *heavily damped* than eyes-closed EC alpha. Such a result would not be compatible with the synchronous oscillator model (Fig 1A) but rather the damped oscillator model (Fig.1B).
2. Cortical input will be significantly different between EC and EO conditions based on the numerous electrophysiological and functional neuroimaging studies that posit a central role for thalamo-cortical input in driving cortical alpha-band activity.
3. Ratio changes in alpha damping between EC and EO will be driven by ratio changes in cortical input between EC and EO conditions.
4. NMDA antagonism will alter the relationship between the ratio changes in alpha damping between EC and EO and the ratio changes in cortical input between EC and EO. Specifically ratio changes in cortical input will be associated with proportionally smaller ratio changes in alpha damping.

## 2. Methods and Materials

### 2.1. EEG experiment - participants and procedure

Thirty healthy males aged between 19 and 37 years old participated in the experiment as described in Forsyth et al. (2018). Participants were scanned on three separate occasions in a placebo-controlled, single-blind, three-way cross-over design using ketamine, midazolam, and placebo. Only the ketamine data are used here, as these were necessary and sufficient to answer the stated hypotheses. Racemic ketamine was administered intravenously with a 0.25mg/kg bolus dose, followed by a 0.25mg/kg/hr infusion. The experimental procedure relevant here commenced with a two minute eyes closed period. Sixteen minute resting state scans were then obtained with drug administration commencing seven minutes into the scan with participants instructed to have their eyes open and fixated on a small cross on a projection screen. This was then followed by a two minute eyes-closed period. For the eyes-open data we used the first and last two minutes of the eyes-open scan for the pre and post eyes-open data respectively. Hence, we had two minute periods with both eyes open and eyes closed for pre and post ketamine infusion available for analysis. The data used to generate the results in this manuscript will be made available on request to Dr Muthukumaraswamy. Access to the data will not be unreasonably withheld.

### 2.2. EEG experiment - data acquisition and preprocessing

EEG data was recorded continuously using Brain Products (Brain Products GmbH) equipment; two BrainAmp MR plus amplifiers with 64-channel Braincaps. Electrode caps used the manufacturer standard layout as described in Forsyth et al. (2018). Data were recorded with a sampling rate of 5 KHz with filters of 0.1-250 Hz with electrode impedances below 10 kΩ and referenced to a common average. Offline, following gradient artefact correction (Allen et al., 2000), data were down-sampled to 500 Hz, with a 100 Hz low-pass filter applied and ballistocardiogram artefact removed (Liu et al., 2012). Visual inspection was used to manually remove further large artefacts and then ICA used to remove other artefact components (e.g. blinks). Data from three participants were excluded at this stage due to excessive artefacts.

### 2.3. Liley mean field model

To model the shape of the resting EEG spectrum in eyes-closed and eyes-open conditions, and how it changes in response to pharmacological manipulation, we utilise the well-known electrocortical mean field model of Liley et al (Liley, 2014; Liley et al., 2002). Mean field models, which describe the average dynamics of populations of neurons, are well suited to describing the EEG, as for a single recording electrode the recorded activity represents the summed activity of many thousands of neurons (Murakami and Okada, 2006). Further, because the defining equations are compact, they are capable of yielding analytical and semi-analytical solutions that efficiently enable the systematic exploration of the physiologically admissible parameter space.

The Liley model is constructed at the macro-columnar scale in which locally excitatory and inhibitory neurons interact by all possible combinations of short-range axo-synaptic feed-forward and feed-back intra-cortical connectivity. This model can be extended to the whole brain scale by allowing macrocolumns to interact via long range excitatory cortico-cortical fibres, however, such an inclusion does not change the essential form of any subsequent linearisation (see below). Based on this macrocolumnar topology dynamical equations of motion for the mean soma membrane potential of excitatory (*h_e_*) and inhibitory *(h_i_*) neurons, averaged over the scale of the macrocolumn, are formulated assuming a simple conductance-based mean neuron. The connection with electrophysiological experiment/measurement is through *h_e_*, which is assumed to be linearly related to the EEG. Mathematically the model can be written as the following coupled set of non-linear differential equations:

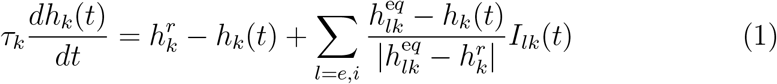

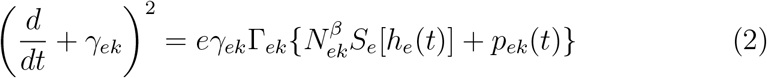

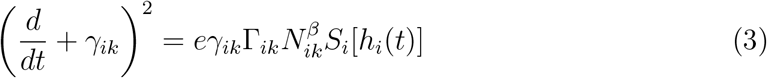

where *k,l* ∈ {*e*(xcitatory), *i*(nhibitory)} and 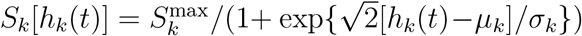 is a sigmoidal function relating the mean soma membrane potential *h*_k_ to the corresponding neuronal population firing rate *S_k_*. Equation (1) represents a single compartment neuron into which all synaptic currents terminate. Equations (2) and (3) describe the dynamics of the excitatory and inhibitory postsynaptic potentials in response to the respective afferent spike rates *S_k_* [*h_k_* (*t*)]. For a definition of all model parameters see Table 1 in Appendix A.

Numerical simulations of these non-linear equations, and various extensions, reveal a wide range of dynamics - which include alpha-band limit cycle activity, multistability and parametrically extensive chaos (Dafilis et al., 2015, 2013, 2009; van Veen and Liley, 2006; Liley et al., 2002; Dafilis et al., 2001). However, linearisations of these equations reveal many important physiologically relevant features in particular electroencephalographically plausible spectra to white noise driving for a wide range of physiologically admissible parameterisations (Hartoyo et al., 2019; Bojak and Liley, 2005; Liley et al., 2002). Such a linearisation is further justified by the empirical results of non-linear time series analysis in which it has been demonstrated that resting EEG, and EEG recorded during anaesthesia, are found to be statistically indistinguishable from a linearly filtered random process (Jeleazcov et al., 2006; Stam, 2005; Jeleazcov et al., 2005; Jeleazcov and Schwilden, 2003; Schwilden and Jeleazcov, 2002; Stam et al., 1999). Linearising Equations (1-3) about a steady state 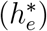 and assuming *γ_lk_* ≡ *γ_l_* (i.e. postsynaptic potential kinetics depend only on the presynaptic source), yields in the frequency domain (*ω*) the following expression (Liley et al., 2002):

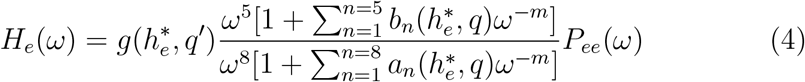

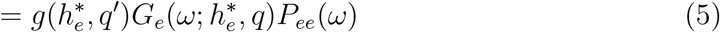

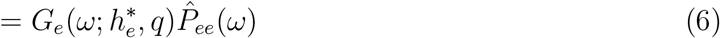

where it has been assumed that only fluctuations in excitatory input to the excitatory cortical neuronal population are non-zero. *H_e_*(*ω*) and *P_ee_*(*ω*) are the Fourier Transforms of *h_e_*(*t*) and *p_ee_*(*t*) (excitatory cortical input on excitatory cortical neurons) respectively. The vector *q* corresponds to the model’s parameters (see Table 1 in Appendix A), and 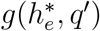 represents a gain parameter that depends on a subset of these parameters *q′* ∈ *q*. 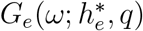 is typically referred to as the *Electrocortical Transfer Function*. The coefficients *a_n_* and *b_n_* will in general be complicated expressions involving 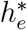 and *q*, the exact form of which is not relevant to the form of 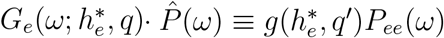 and can be understood as the frequency domain representation of afferent cortical input/driving.

Such a linearisation, parameterised by physiologically admissible values, is able to well account for the shape of the resting eyes-closed and eyes-opened EEG spectrum (Hartoyo et al., 2019; Bojak and Liley, 2005; Liley et al., 2002). Figure (2) shows some representative comparisons between empirical power spectral densities (PSDs) and PSDs estimated from |*H_e_*(*ω*)|^2^, on the basis of maximum likelihood estimation using a Markov chain Monte Carlo method. By systematically calculating the sensitivity of the estimated PSD shape to parameter variations it is found that parameters *γ_i_* (the IPSP shape/decay) and *γ_e_* (the EPSP shape/decay) play the dominant role in determining PSD shape. In particular it is found that *γ_i_* principally determines centre alpha frequency, whereas *γ_e_* dominantly determines alpha peak width (or full-width half maximum, FWHM) (Hartoyo et al., 2019).

**Figure 2:**
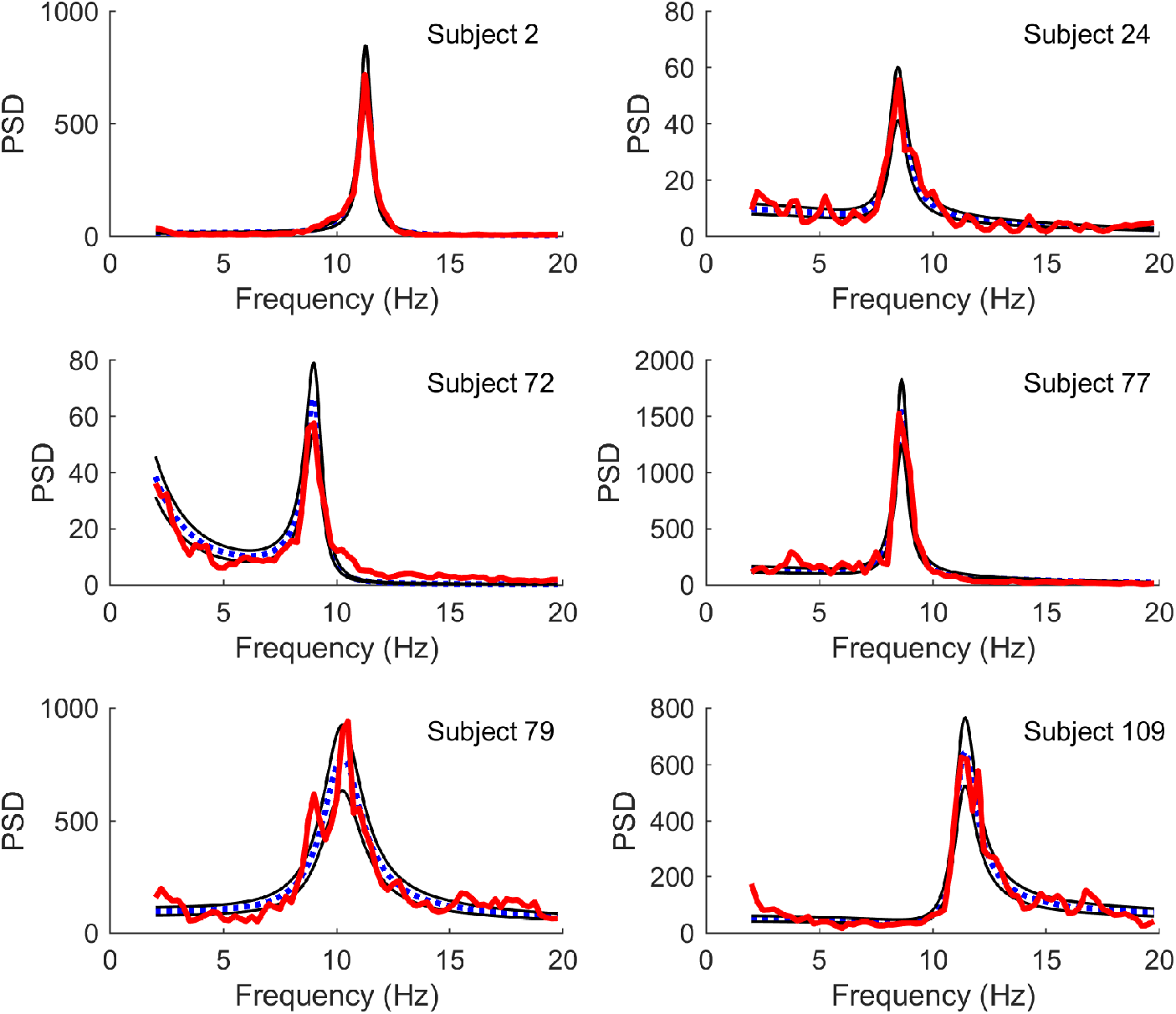
Comparison of model PSDs (Equation 4, blue dotted line) fit to experimental PSDs (red thick line) by a maximum likelihood estimation using a Markov chain Monte Carlo method for a number of subjects. Also shown are the 16% and 84% quantiles based on the gamma distribution for the fitted spectra (thin black lines). Experimental spectra were calculated in the resting eyes-closed condition. For further details see Hartoyo et al. (2019). Figure reproduced from Figure 1 of Hartoyo et al. (2019)

### 2.4. Time series analysis

The model linearisation defined by Equation (4), based on assuming a pole-zero matching, leads to a convenient fixed-order Auto-Regressive Moving-Average (ARMA) process which can be used as a parsimonious model for the analysis of recorded EEG. The fixed-order ARMA model so obtained is of autoregressive (AR) order 8 and moving average (MA) order 5. This can be easily understood from Equation (4) in which the differential operator in continuous time (i.e. *d/dt* → *iω*) becomes a difference operator in discrete time, such that the 5th and 8th order temporal derivatives in the numerator and denominator of *G_e_* become respectively 5th and 8th order finite differences in discrete time. These theoretically derived fixed AR and MA orders accord well with empirical determinations of optimal AR (range 3–14) and MA (range 2–5) orders obtained from resting awake eyes-closed EEG using a range of information theoretic criteria (Schack & Krause 1995; Tseng et al 1995). Such a fixed order ARMA time series method has been used successfully to differentiate the effects that a variety of pharmacologically diverse agents have on resting MEG/EEG/ECoG recordings (Muthukumaraswamy and Liley, 2018; Shoushtarian et al., 2016b,a; Liley et al., 2010, 2008, 2003).

Therefore, for a discretely sampled EEG time series, *y*[*n*], we posit that it can, for stationary intervals, be described in terms of the following difference equation

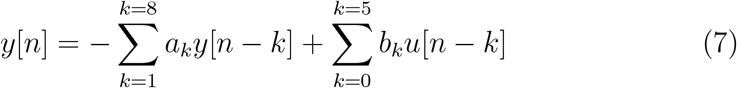

where *a_k_*, *b_k_* are the AR and MA coefficients respectively, *b*_0_ = 1 and *u*[*n*] is an iid sequence of random variables having variance 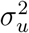. By defining the Z-transform of a discrete time sequence *x*[*n*] as 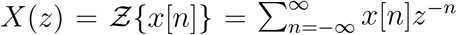 this can rewritten as

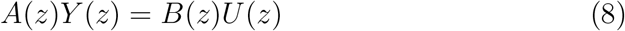

where *A*(*z*) and *B*(*z*) are the following polynomials *A*(*z*) = 1 + *a*_1_*z*^-1^ + ⋯ + *a*_8_*z*^-8^, *B*(*z*) = 1 + *b*_1_*z*^-1^ + ⋯ + *b*_5_*z*^-5^. Solutions to *A*(*z*) = 0 give the system (filter) poles whereas solutions to *B*(*z*) = 0 give system (filter) zeros. An estimate of the variance of the innovating random process can be obtained as 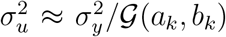, where is the variance of the recorded signal and 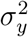 is the *power gain* of the estimated filter to a unit variance white noise driving process. As in previous publications we will refer to *σ_u_* as the *cortical input* (CI) (Liley et al., 2010). The estimated AR and MA coefficients define a power spectral density function according to

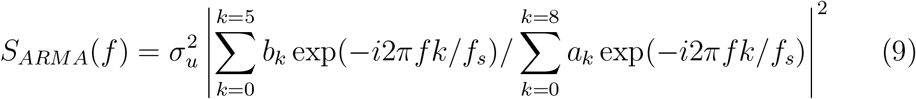

It is of note that such time series models are often preferred for spectral estimation of stochastic processes when we have *a priori* knowledge regarding the underlying generative mechanism (Broersen, 2002).

The poles 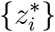 that are solutions to *A*(*z*) = 0 can be thought of as the dominant system oscillatory modes that when stochastically perturbed and summed approximate the recorded time series *y*[*n*]. Each pole 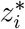 has a frequency 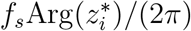 and a damping 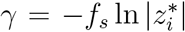, where *f_s_* is the sampling frequency. The damping parameter quantifies how rapidly an oscillation of frequency 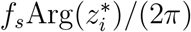 decays over time after it has been perturbed/excited. For a Lorentzian power spectral density FWHM = 2*γ*.

In order to calculate the ARMA coefficients {*a_k_, b_k_*} we down-sampled data to 80 Hz and fitted an ARMA (8,5) model to the data using the AR-MASA MATLAB toolbox (Broersen, 2002) to 2 s 50% overlapping segments. We then selected ‘alpha’ poles having frequencies lying between 5 and 15 Hz in order to calculate the corresponding damping. For topographic plots the median dampings of alpha poles, and the median CI, for each EO or EC condition were used.

In order to clearly differentiate changes in alpha power that are driven by changes in cortical driving and changes in the damping of cortical alpha we plot the ratio change in alpha power calculated by our fixed order ARMA versus the ratio change in the estimated variance of the white noise driving process between EC and EO conditions. To do this we assume that our ARMA calculated alpha power depends on the integral of a Lorentzian distribution parametrised by the estimated pole damping *γ* and frequency *f_α_*,

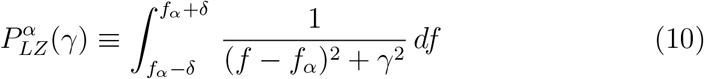

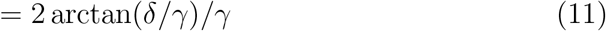

where *δ* = 2.5 Hz. Such a calculation allows us to estimate changes in alpha-band spectral power that are driven by changes in alpha pole damping and not from the extended spectral effects of any of the other poles that contribute to the overall power spectral density as estimated by Equation (9). Total alpha power is then estimated as 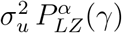. Non-parametric frequency analysis was performed using the FieldTrip toolbox (Oostenveld et al., 2011). Specifically the power spectral density was calculated using multiple tapers based on discrete prolate spheroidal sequences with smoothing of 2 Hz.

## 3. Results

Thirty healthy participants had simultaneous EEG-fMRI recordings before and during the intravenous administration of the putative NMDA-receptor antagonist ketamine. Of these twenty seven had EEG data suitable for further analysis. Comparisons between the power spectral densities between EC and EO conditions calculated using a theoretically-motivated fixed order autoregressive moving-average (ARMA) model of MA order 5 and AR order of 8 (see Time Series Analysis in Materials and Methods) and FFT are illustrated in Figures 3A & B for electrode PO4 (site of maximum group-level change - see 4B). The strong similarity in the shape of the estimates of the power spectral density and the similarity in differences in alpha-band attenuation are clearly evident, with the only major difference being a low frequency peak arising from the high-pass filtering necessary for the calculation of the FFT power spectral estimate. Figure 3C illustrates for one participant the clear increase in the full-width half-maximum (FWHM) during the EO condition indicating (see Fig. 1B) that the alpha oscillatory mode is more heavily damped. However, in general, because of the co-existence of significant delta band (0–4 Hz), the FWHM, and hence the damping of the alpha oscillatory mode, cannot be directly evaluated from estimates of power spectral density (Fig. 3D, also see Fig. S1 in SI). For this reason fixed order ARMA methods need to be used in which alpha oscillatory mode dampings’ can be more reliably estimated.

**Figure 3:**
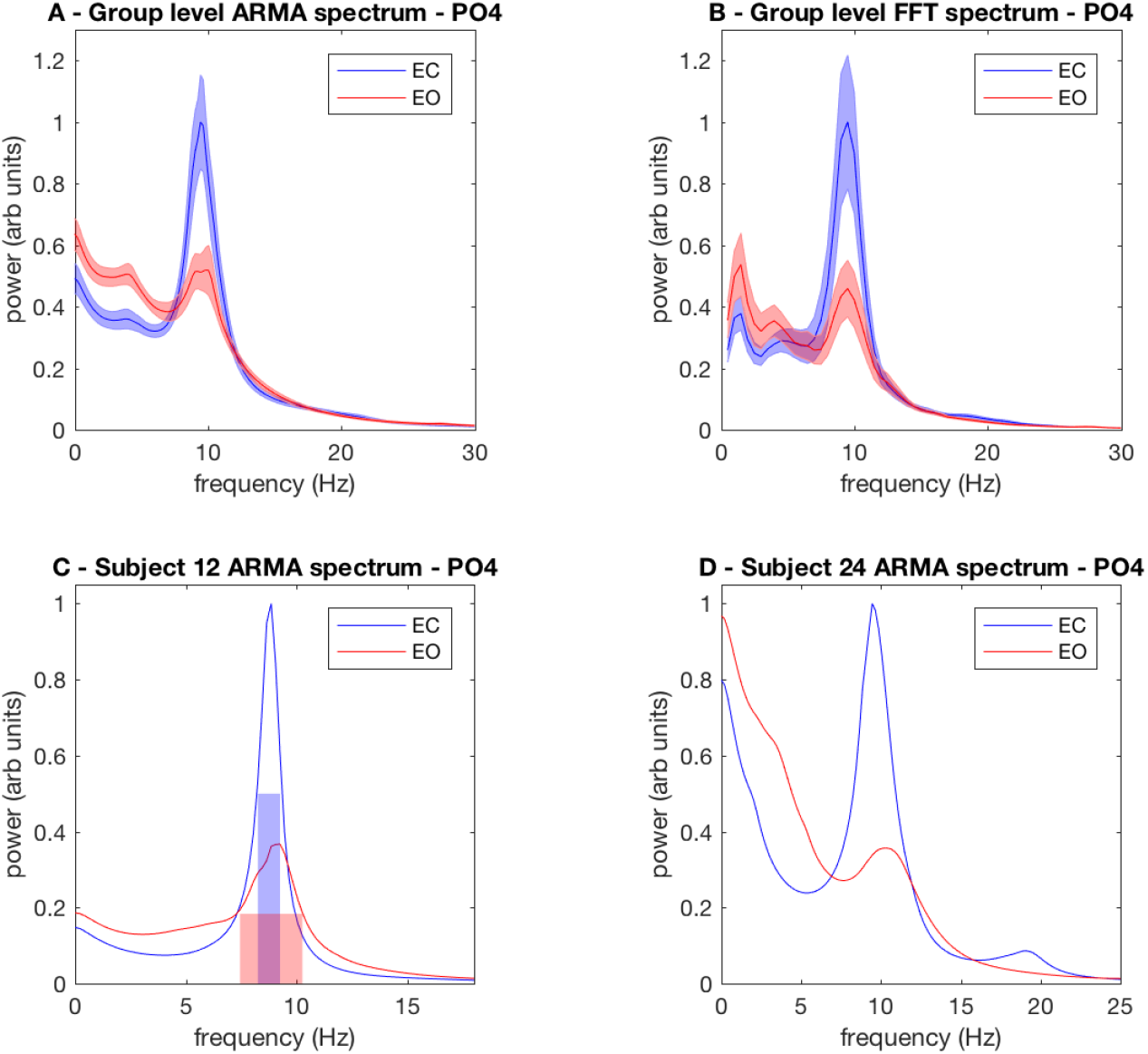
Comparisons between EC and EO conditions for FFT and ARMA calculated power spectral density functions. A - Group-level fixed order ARMA power spectral density estimates for EC and EO conditions at the electrode associated with maximal group-level alpha oscillatory mode damping (see Fig. 4B); shaded areas represent standard error of the mean. B - Group-level multitaper FFT power spectral density estimates for EC and EO conditions at same electrode location as A. C - Illustration of a case in which the changes in alpha FWHM, which will be proportional to the change in the damping of the corresponding alpha oscillatory mode, can be evaluated directly from the power spectral density estimate (ARMA spectral density illustrated only). D - Illustration of a case where changes in the alpha FWHM cannot be directly evaluated from the power spectral density estimate.

As expected we find a strong relationship between non-parametric multitaper FFT and parametric fixed order ARMA(8,5) spectral estimates of EC-EO changes in alpha-band (8–13 Hz) power (Fig. S2A in SI). A clearer understanding of the basis for this relationship is obtained by assuming that the parametrically calculated differences in alpha-band power depend only upon the changes in damping of a single alpha oscillatory mode. In particular, we find that the changes in FFT calculated alpha power between EC and EO conditions are well accounted for by the differences in a Lorentzian power spectral density function parametrised by the respective EC and EO dampings of a single alpha oscillatory mode (see Eqs. 10 & 11 in Materials and Methods and Fig. S2B in SI). The Lorentzian power spectral density function (integrand of Eq. 10) is the simplest and most general form for the spectral resonance of a simple damped oscillatory process.

Figure 4A shows group-level topographic changes in alpha-band (8–13 Hz) power between EO and EC conditions during the baseline condition before the commencement of an intravenous ketamine infusion. Most participants exhibited the classically described attenuation of occipital alpha-band power following eyes-opening, with a small number (5-6) not showing any clear effect (see Fig. S2 in SI). In contrast, changes in the damping of the stochastic alpha oscillatory component, calculated using the fixed order ARMA model reveal topographically much more differentiated and widespread increases between EC and EO (Fig. 4B, Fig. S4 in SI). Of particular note is that the locations of increases in damping display only partial overlap with decreases in power. Significantly there are also topographically widespread reductions in the frequency of the stochastic alpha oscillatory component between EC and EO conditions (Fig. 4C, Fig. S5 in SI). Significant changes in the amplitude of the estimated white noise driving process are however restricted to only a few electrodes (Fig. 4D, Fig. S6 in SI).

**Figure 4:**
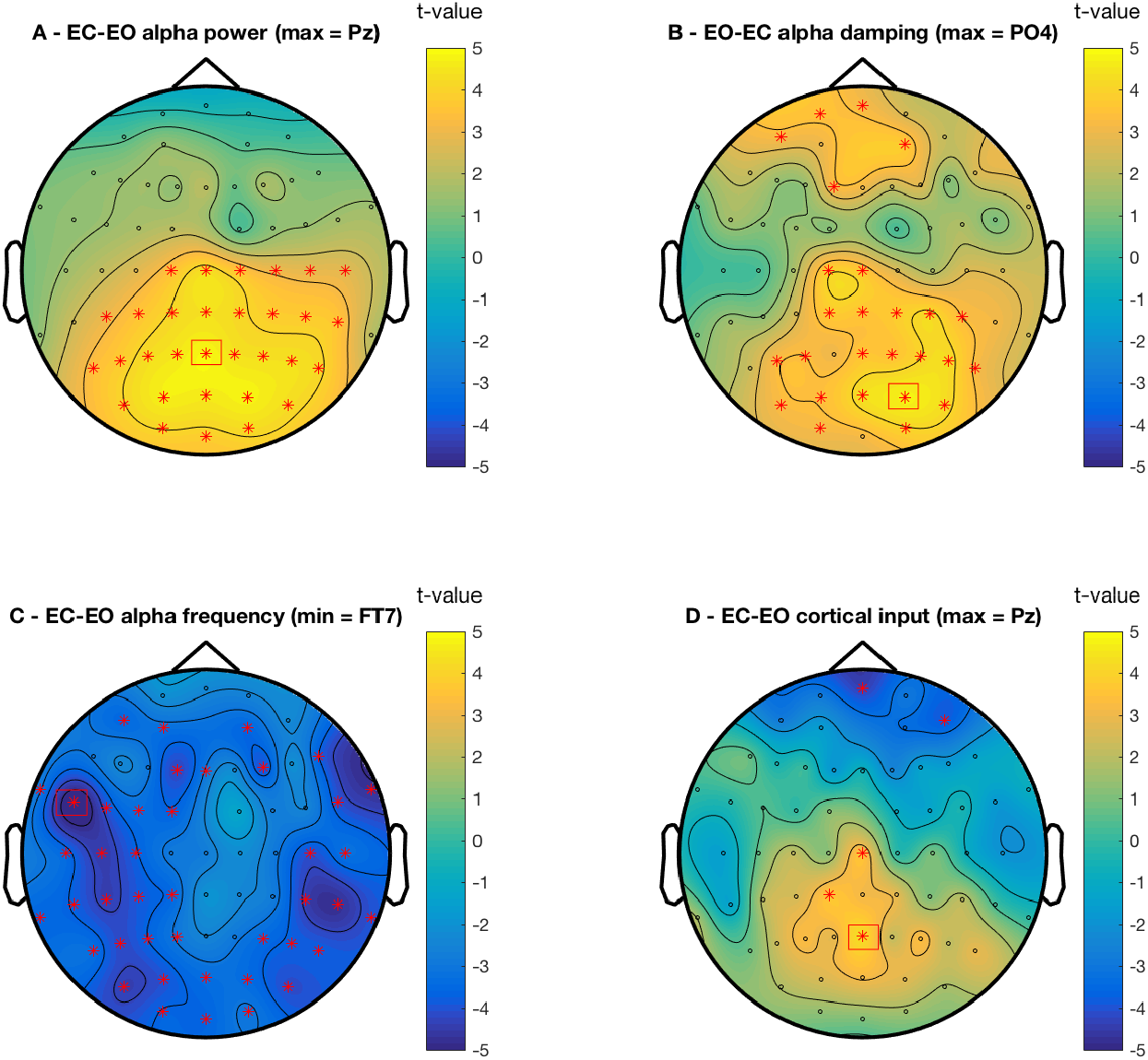
Group-level changes between EC and EO conditions in the baseline condition plotted as topographic maps of t-values. A - EC-EO alpha-band (8-13 Hz) power changes. B - EO-EC alpha oscillatory mode damping changes. As the damping in EO will be greater than in EC we have chosen to plot the EO-EC damping to more easily facilitate topographic comparisons with the corresponding alpha-band power changes. C - EC-EO alpha oscillatory mode frequency changes. D - EC-EO cortical input. Red asterisks represent group-level t-values thresholded at a false discovery rate of 0.05 calculated using a two- level nonparametric permutation test (Nichols and Holmes, 2002). Red boxes represent the electrode location of extremal values.

Figure 5 illustrates, for a single typical participant, the changes seen in the distributions of the estimated damping, frequency and the amplitude of cortical input between EO and EC conditions. A small number of participants however exhibited little change in the respective distributions whereas others evinced clear changes in either the damping or the amplitude of cortical input (see Fig. S7 in SI). A statistical analysis across all participants (Fig. 6 - top row) for changes in alpha oscillatory mode damping and frequency, cortical input and FFT calculated alpha-band (8–13 Hz) power at the electrode location that is associated with the greatest damping between EC and EO conditions reveals only a significant difference in the damping. Because such a result may have been biased by the choice of electrode we chose to aggregate all epochs, participants and electrodes in order to form group-level distributions for damping, frequency, driving amplitude and alpha (8–13 Hz) power in EO and EC conditions (Fig. 6 - bottom row). Non-parametric tests and visual inspection reveal unequivocal differences between the respective distributions for alpha oscillatory mode damping and frequency between EC and EO conditions. As hypothesised alpha oscillatory mode damping and frequency clearly increase between EC and EO states. Less clear, but nevertheless significant, differences are seen in alpha-band power, whereas no significant difference is found between the respective distributions of the magnitude of cortical input i.e. the amplitude of the innovating (driving) process. We find that the ratio change between EC and EO FFT estimated power (EC/EO = 1.22) is similar to the ratio change between EC and EO power calculated between 8–13 Hz using the Lorentzian approximation of Equation (10) parametrised by the respective median dampings (EC/EO = 1.38) (see SI for further details). That is, the group-level changes in damping are able to predict group-level changes in the non-parametrically calculated FFT alpha-band power to within ~10%. On the basis of the predicted ratio change in EC/EO power it appears that the fixed-order ARMA method can potentially identify latent changes in the oscillatory power of alpha activity that are obscured in the FFT.

**Figure 5:**
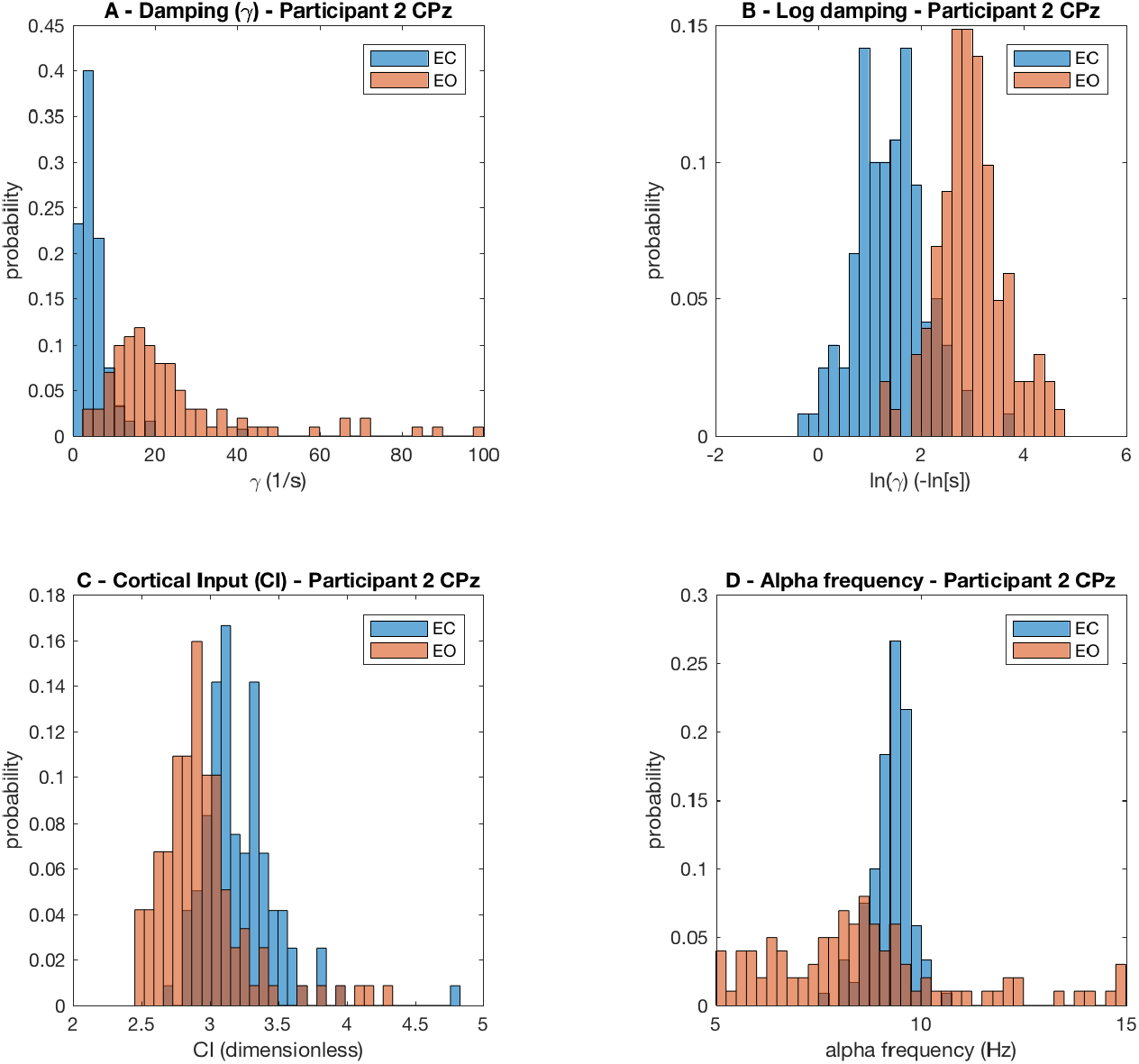
Representative distributions of alpha oscillatory mode damping, frequency and the amplitude of the cortical input (innovating stochastic process (*σ_u_*); see Materials and Methods: Time series analysis) for a typical participant during EC and EO. Most participants exhibited similar changes in manifesting a clear increase in median alpha oscillatory damping.

**Figure 6:**
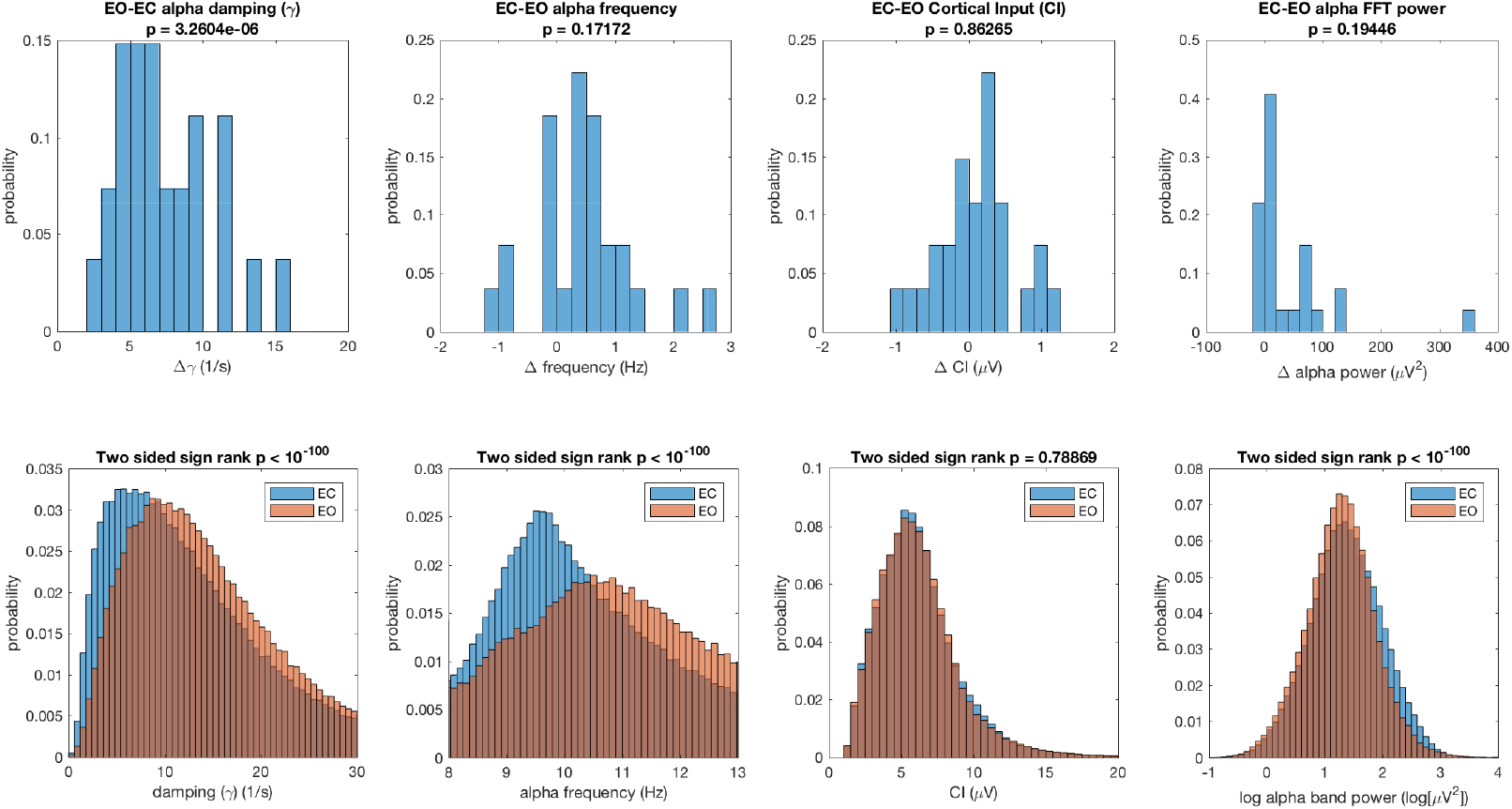
Top row - Group-level changes in maximum damping (and associated frequency and cortical input) and alpha power (8–13 Hz) between EO and EC. Statistical difference from zero mean determined using a two-sided nonparametric sign test. Bottom Row - Histograms of alpha oscillatory mode damping, alpha oscillatory mode frequency, cortical input and log alpha power for all participants, electrodes and epochs for EC and EO. Statistical difference evaluated using unpaired two-sided Wilcoxon rank sum test.

While no significant changes in the amplitude of cortical input between EO and EC states were found overall, this does not preclude the possibility that when evaluated on an epoch by epoch basis there exist correlations between cortical input and damping. We hypothesised not only that ratio changes in alpha damping between EC and EO will be driven by ratio changes in cortical input between EC and EO, but also that this relationship will be significantly altered following NMDA antagonism by the putative NMDA antagonist ketamine. Figure 7-top row illustrates ratio changes in ARMA calculated alpha power and differences in alpha oscillatory mode damping plotted, between EC and EO states, with respect to ratio changes in the estimated cortical input for both baseline and ketamine conditions. As hypothesised (see Fig. 1C-right panel) we see that in the absence of ketamine there is a strong relationship between cortical input and changes in damping. In contrast, the action of ketamine is to significantly diminish this relationship such that ratio changes in power and differences in damping between EC and EO are essentially independent of ratio changes in the driving input. Such a difference is even more stark when such changes are plotted for the electrode location in each participant in which alpha oscillatory mode damping increased greatest between baseline EC and EO conditions (Fig. 7-bottom row).

**Figure 7:**
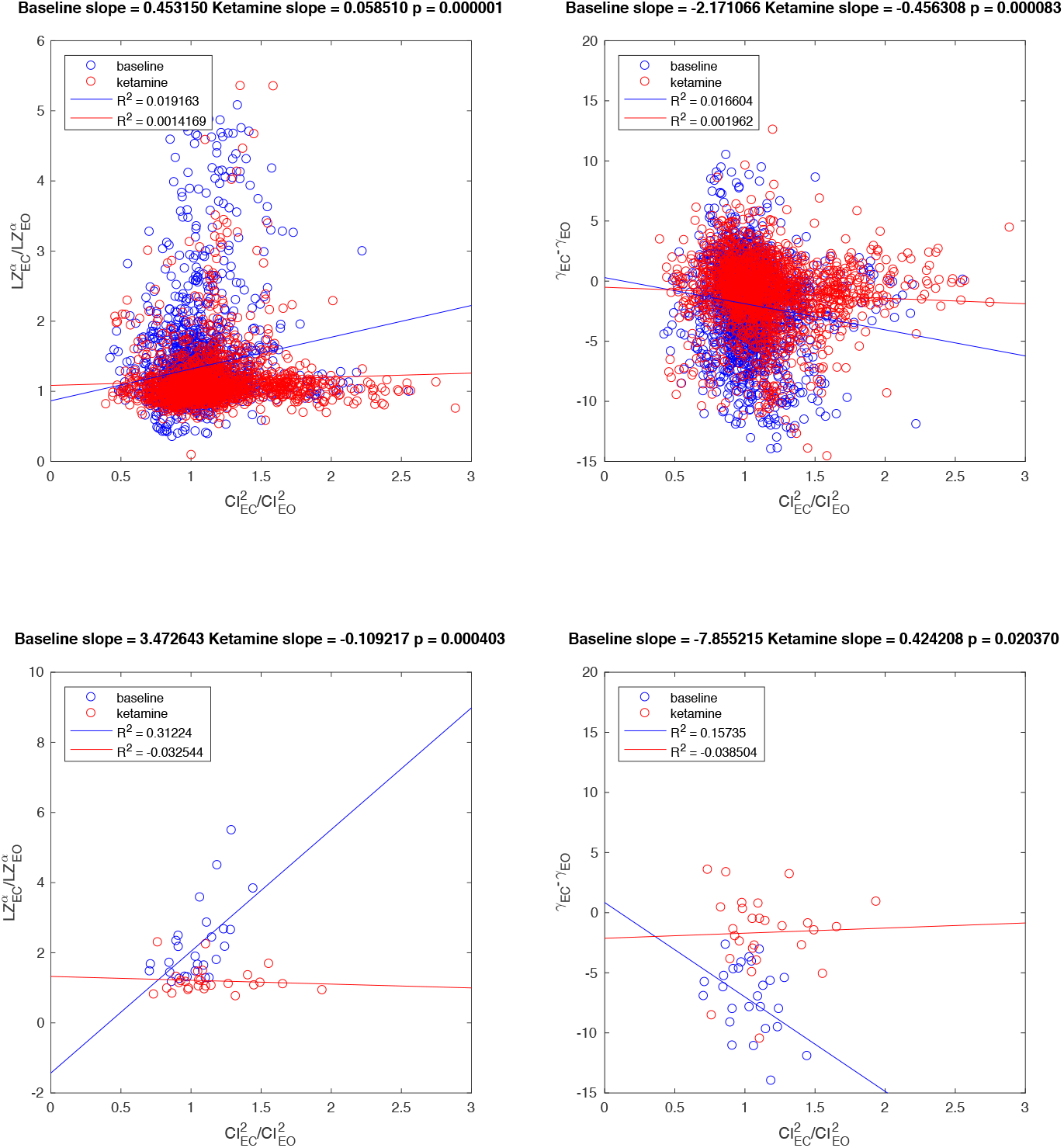
Top row - Ketamine administration in healthy participants modifies the relationship between estimated cortical driving and the ARMA calculated alpha-band power and alpha damping. In the absence of ketamine ratio changes in cortical driving between EC and EO are positively correlated with ratio changes in Lorentzian approximated alpha-band power (left), and with differences in ARMA calculated alpha damping (right). However, following ketamine administration this relationship is significantly diminished such that increases in cortical driving do not appear to drive corresponding increases in alpha oscillatory damping. Results shown are for all participants and all electrodes. Bottom row - Effect of ketamine on alpha blocking for the most heavily damped channel in the baseline (no ketamine) condition. Results should be compared with the right most panel of Figure 1C.

Finally, we investigated whether changes in the damping of alpha could account for the commonly studied event-related alpha amplitude changes. To do this we re-analysed source-level MEG data from a previously published experiment (see MEG experiment - event-related data analysis in SI and (Muthukumaraswamy et al., 2013)). Figure 8 shows the power time-course expressed as a relative change from the baseline amplitude. It can be observed that immediately following stimulus onset (time = 0 s) there is a transient increase in power representing the evoked response followed by sustained alpha amplitude decreases. ARMA-estimated power spectra show a clear reduction in alpha power (Fig. 8B). In Fig. 8C damping estimates for all trials are plotted as histograms with a clear increase in damping seen in the post-stimulus period (median pre = 2.08, median post = 2.41, z= 12.02, p= 2.86e-33). Computing the median for each (Fig. 8D) participant (n=15) and computing a random effects sign rank test showed significantly greater damping post-stimulus (p=1.22e-04).

**Figure 8:**
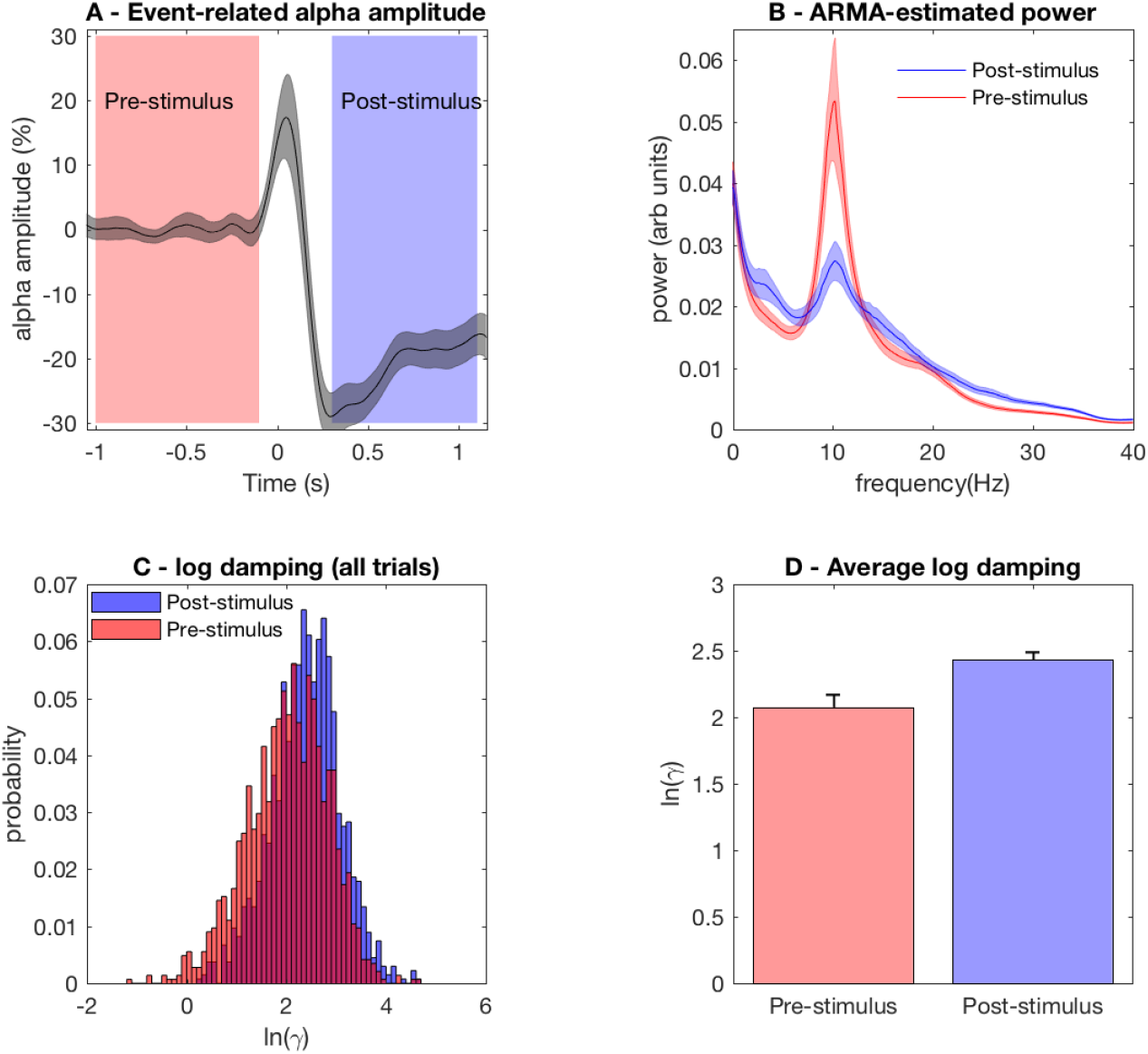
Analysis of damping in source-level MEG data following visual stimulation. A - Time course of oscillatory alpha amplitude expressed as relative change (%) of the baseline period with pre and post stimulus time-windows carried forward in subsequent panels indicated. B - ARMA power spectral density estimates for pre and post stimulus time-windows. C -Probability of oscillatory alpha mode damping for each trial and participant. D - Average damping across participants computed by taking the median damping value of each participant.

## 4. Discussion

The standard view regarding the alpha rhythm is that thalamus plays the dominant role in its genesis and modulation (Hughes and Crunelli, 2005; Steriade et al., 1990). In this regard separate explanations have emerged for its generation and the changes in its amplitude that occur in response to a wide range of behaviours and stimuli. For example, the alpha rhythm is typically thought to either arise directly from alpha-band activity generated in the thalamus ‘pacing’ overlying cortical tissue (Hughes and Crunelli, 2005) or due to reverberant activity arising from the reciprocal interactions between both cortex and thalamus (Robinson et al., 2002).

In contrast, the phenomena of alpha blocking, or its event-related counterpart event related desynchronisation (ERD; Pfurtscheller and Lopes da Silva (1999)), is thought to be explicable in terms of a theory in which it was proposed that ‘blocking’ is the result of a reduction in the synchronisation of multiple, more or less identical, alpha oscillators (Elul, 1972). On this basis it seems the terms for describing alpha-band power changes in terms of ‘synchronisation’ and ‘desynchronisation’ (which had existed for some time and were descriptive of the temporal regularity or irregularity of changes in recorded EEG amplitude) became repurposed as a mechanistic explanation. For this reason it is generally believed that mean field theories of the EEG *‘… cannot hope to describe ERD and ERS [event related synchronisation], at least not at the single population level’* (Coombes and Byrne, 2019) because they define the temporal evolution of neuronal population dynamics and therefore make an implicit assumption of synchrony. Because of the common belief that brain dynamics will only be explicable in terms of the behaviour of individual neurons the idea that amplitude changes observed in the EEG are the result of desynchronising/synchronising factors has therefore, understandably, prevailed. For example Nunez and Srinivasan (Nunez and Srinivasan, 2005) note that *“The large changes in scalp amplitude that occur when brain state changes are believed to be due mostly to such synchrony changes”*.

However, we have here presented macroscopic evidence that provides a definitive challenge to such a ‘microscopic’ explanation for a bulk electrocortical phenomena. If alpha blocking was the result of the desynchronisation of a population of near identical oscillators we would not expect *any* change in the relative distribution of power within the alpha-band. From our systems-level perspective we would therefore expect the damping of alpha to remain unaffected but the amplitude of the driving white noise process (cortical input) to decrease. Statistically we do not observe any significant reduction in cortical input and as such it cannot account for the observed changes seen in alpha power between EC and EO conditions. Instead we find that alpha is more heavily damped in the EO condition compared to EC. In other words the change seen in alpha power is not the result of ‘microscopic’ level desynchronising processes but the result of larger-scale macroscopic alterations in the system properties of cortical neuronal population activity. Additionally we find that the median frequency of the alpha resonance significantly increases between EC and EO conditions (Fig. 6 bottom row) mirroring recent results in which it is argued that increases in alpha frequency reflect reductions in cortical excitability (Samuel et al., 2018) or alterations in cognitive demand (Haegens et al., 2014). Such combined changes in alpha frequency and damping can form the basis for parametric explorations in mean field models in order to better understand their physiological genesis (Liley et al., 2002). However, while we find no direct evidence for changes in sub-cortical/thalamic input driving changes in alpha-band amplitude we cannot discount the possibility that thalamus is having a modulatory role on cortical dynamics (Sherman and Guillery, 2005). Indeed our approach cannot exclude the possibility that thalamic, and cortico-cortical, input are driving the observed changes in alpha-band damping between eyes-closed and eyes-open conditions, as we discuss further below.

In this work we have chosen not to investigate whether changes in the strength of coupling of a population of phase oscillators could mechanistically account for the observed changes in the shape of the alpha-band power spectrum. Such an approach, typically investigated using the Kuramoto model (Acebron et al., 2005), assumes that a population of weakly-coupled limit cycle oscillators, having a distribution of natural frequencies, are able to progressively synchronise as the coupling strength is increased above a certain critical threshold. While this model has found widespread use in the neurosciences (Breakspear et al., 2010; Cumin and Unsworth, 2007) its application here would be problematic. Firstly, and most significantly its dynamics are not able to account for the shape of the alpha-band resonance. In the frequency domain changes in synchronisation in the context of the Kuramoto model are revealed as changes in the amplitude of a single frequency, necessarily reflecting the entrainment of limit cycle activity. But as previously noted (Methods - Liley mean field model) there is scant evidence to support the notion that resting EEG is indistinguishable from a filtered random process. Secondly, changes in the magnitude of synchronisation in the context of the Kuramoto model (in the thermodynamic limit) will not be associated with any change in the entrained frequency. In contrast we have identified clear changes in alpha-band frequency between EC and EO conditions. Finally, the Kuramoto model is a mathematical model of synchronisation only, it tells us little about the dynamical and physiological mechanisms responsible for the genesis of the oscillations and as such cannot be used to meaningfully link the underlying physiology to any emergent dynamics.

By using ARMA methods we have been able to first-order identify changes in cortical input that are the consequence of changes in EO and EC states. In the normal state we observe a clear correlation between cortical input and the damping of alpha oscillatory modes, which at first sight might be seen to support the role of the thalamus in the genesis of the alpha rhythm. However, thalamo-cortical connections only contribute of the order of 5% of all afferent connections to a given region of cortex (Schoonover et al., 2014; Braitenberg and Schüz, 1998), with the vast majority arising from cortico-cortical inputs from other areas of cortex. Thus it is not only thalamic input to cortex that will be expected to drive changes in the damping of alpha. We had hypothesised that increases in cortical input would drive neuronal population activity and by doing so would increase the contributions of voltage dependent NMDA receptor mediated ionic currents to unitary excitatory post-synaptic currents by prolonging their decay phase (see Fig. 1C). Theoretically the predicted effect of this would be to increase the amplitude of the alpha rhythm by reducing its damping. On this basis we were able to reasonably predict that blocking the effects of NMDA receptor action in cortex should be associated with a reduction in the effect that changes in cortical input will have in driving changes in the damping of alpha-band oscillatory activity. Indeed this is what we found following the administration of ketamine to healthy participants (Fig. 7).

In conclusion we have presented a clear argument, supported by definitive data, that the alpha rhythm and its blocking can be explained by a single, physiologically well-motivated, theory. We have therefore effected a reduction between a theory regarding the genesis of the alpha rhythm and a theory for its attenuation to a single well defined explanation. Therefore, use of the terms ‘synchronisation’ and ‘desynchronisation’ may be seen as misleading in terms of mechanistically explaining changes in large scale alpha-band power. On the basis of the results of Fig. 8 it would seem the terms Event Related Desynchronisation (ERD) and Event Related Synchronisation (ERS) may need to be replaced with the mechanistically more informative terms Event Related Damping (‘ERD’) and Event Related Amplification (ERA), at least in the context of alpha-band oscillatory activity.

## Acknowledgments

We thank Anna Forsyth and Rebecca McMillan (University of Auckland) for help in data collection. Juergen Dukart and Jörg Hipp (Roche Pharma Research and Early Development, Neuroscience, Ophthalmology and Rare Diseases, Roche Innovation Center, Basel, Switzerland) kindly advised on aspects of data pre-processing and collection. The data analysed in this paper was funded by a grant from F Hoffman La Roche Ltd.

## Appendix A. Liley mean field model parameters

**Table A.1:** Model parameter definitions for Equations (1 – 3) that mathematically represent the macrocolumnar Liley model (Liley, 2014; Liley et al., 2002)

| Model parameter | Definition |
| --- | --- |
| $h_e^r, h_i^r$ | resting membrane potential of excitatory, inhibitory neurons |
| $h_{ee}^{\text{eq}}, h_{ei}^{\text{eq}}$ | excitatory reversal potential in excitatory, inhibitory neurons |
| $h_{ie}^{\text{eq}}, h_{ii}^{\text{eq}}$ | inhibitory reversal potential in excitatory, inhibitory neurons |
| $\tau_e, \tau_i$ | passive membrane time constant for excitatory, inhibitory neurons |
| $\gamma_{ee}, \gamma_{ei}$ | EPSP rate (shape/decay) constant in excitatory, inhibitory neurons |
| $\gamma_{ie}, \gamma_{ii}$ | IPSP rate (shape/decay) constant in excitatory, inhibitory neurons |
| $\Gamma_{ee}, \Gamma_{ei}$ | EPSP amplitude in excitatory, inhibitory neurons |
| $\Gamma_{ie}, \Gamma_{ii}$ | IPSP amplitude in excitatory, inhibitory neurons |
| $N_{ee}^\beta, N_{ei}^\beta$ | total number of excitatory synapses per excitatory, inhibitory neuron |
| $N_{ie}^\beta, N_{ii}^\beta$ | total number of inhibitory synapses per excitatory, inhibitory neuron |
| $S_e^{\text{max}}, S_i^{\text{max}}$ | maximum mean firing rate of excitatory, inhibitory neurons |
| $\mu_e, \mu_i$ | mean firing thresholds of excitatory, inhibitory neurons |
| $\sigma_e, \sigma_i$ | std. dev. of firing thresholds of excitatory, inhibitory neurons |
| $p_{ee}, p_{ei}$ | excitatory cortical input in excitatory, inhibitory neurons |

## Supplementary Information

### Ratio EC-EO power calculation

Based on the median damping and frequency of the estimated alpha oscillatory mode over all electrodes, subjects and participants for EC and EO conditions we can predict the corresponding ratio change in non-parametrically calculated Fourier power by using a Lorentzian approximation. Alpha-band spectral power is estimated from the median damping and frequency of the ARMA calculated alpha oscillatory mode as

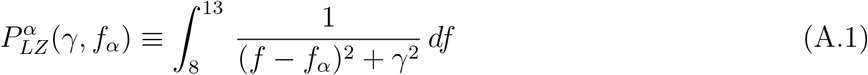

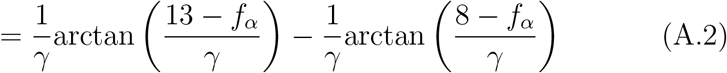

On this basis, because cortical input is not significantly different between conditions (see Fig. 5 in main text), the ratio change in alpha-band power between EC and EO conditions is estimated as

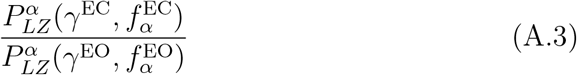

where 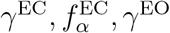 and 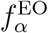 are the respective median dampings and frequencies, in units of *s*^-1^, of the alpha oscillatory mode in EC and EO conditions. For 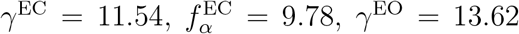 and 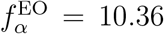 (estimated from the respective histograms in the bottom row of Fig. 5 in main text) this ratio is calculated to be 1.38. The corresponding EC/EO ratio in FFT calculated power is 1.22.

### MEG experiment - event-related data analysis

Data from fifteen participants from a previously published MEG experiment were re-analysed using data from the placebo baseline period (see Muthukumaraswamy et al. (2013) for details). Briefly, the paradigm consisted of 120 trials of 1-1.5s of visual stimulation with a grating patch followed by a manual response and then a 1.5s baseline period. The source-level virtual electrodes sensitive to alpha power changes were used - leaving one sensor per participant to analyse (Muthukumaraswamy et al., 2013). In order to obtain the time series of amplitude changes a 4 Hz wide Butterworth bandpass filter was used centered at 10Hz and amplitude time series obtained using the Hilbert transformation. The pre-stimulus period was defined as −1 to −0.1 s and the post-stimulus period as 0.3 −1.1s. ARMA spectral analysis followed as described above.

**Figure S1:**
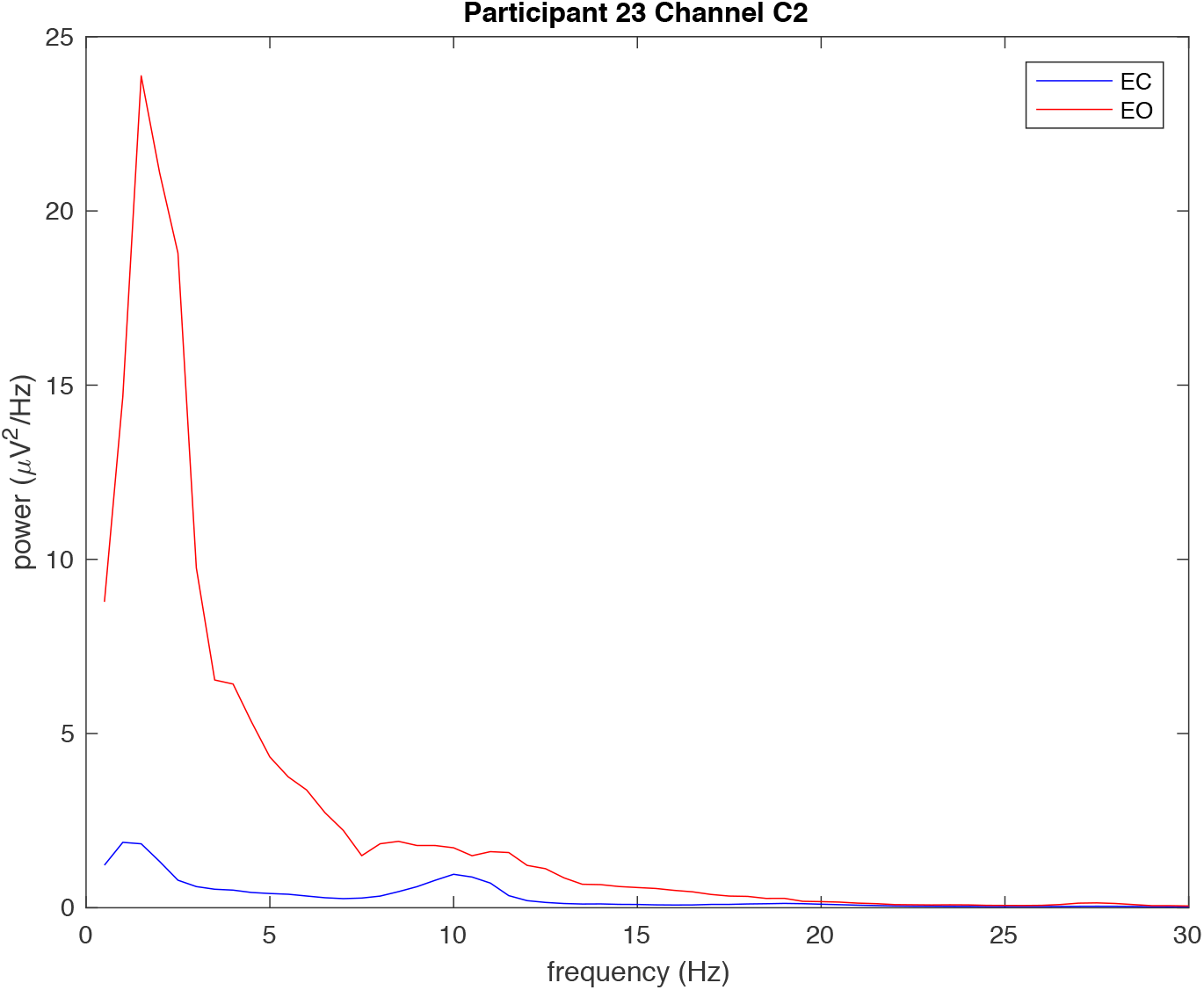
EO and EC FFT power for Participant 23 electrode C2 (red asterix in Figs. S3-S6). Even though judged by FFT analysis this site does not indicate EC-EO blocking ARMA analysis confirms that the damping of the alpha pole has been increased during EO. In this example it appears low frequency oscillatory activity has increased the apparent alpha power thus occluding any assessment of alpha blocking.

**Figure S2:**
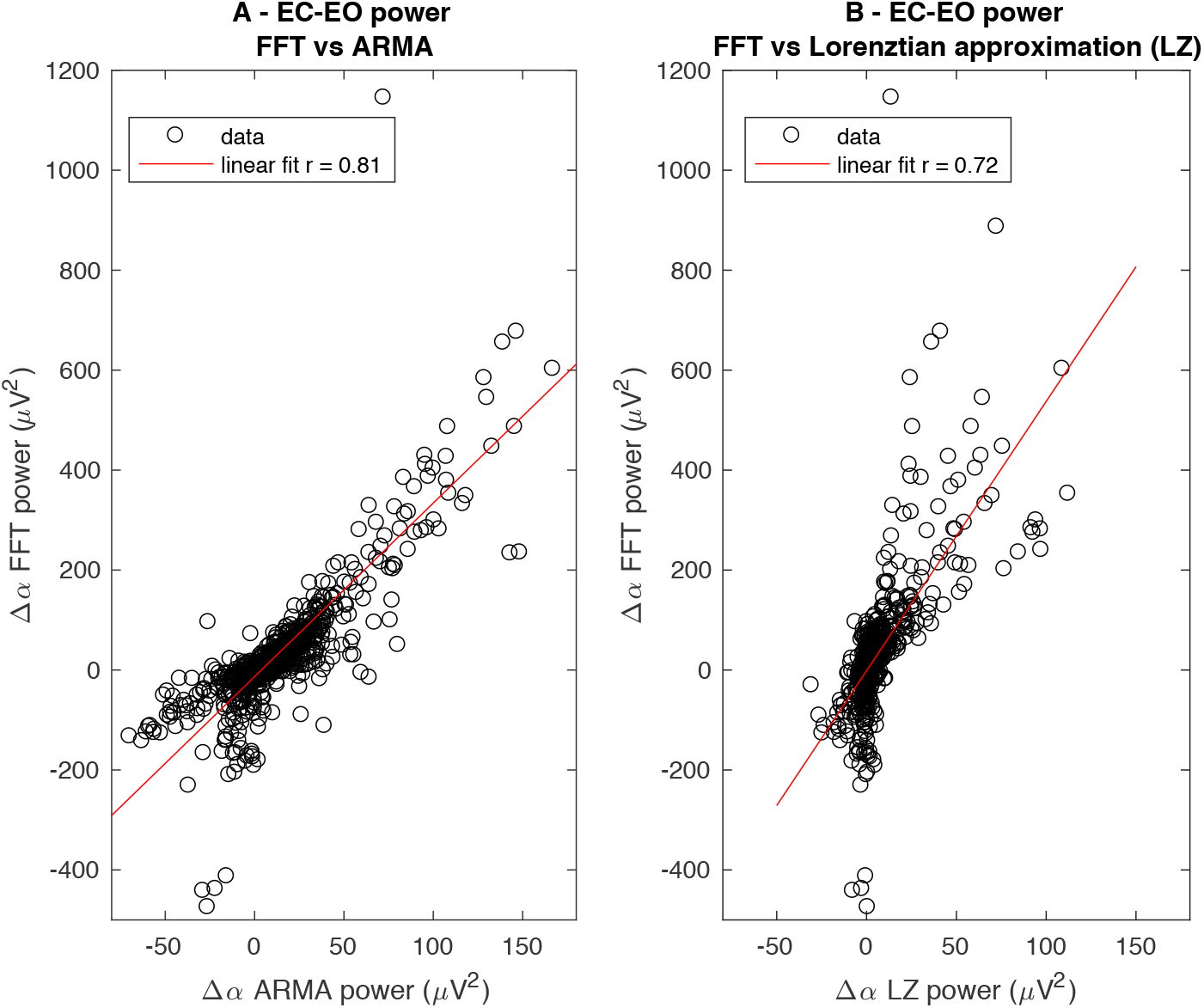
Relationship between changes in non-parametrically and parametrically estimated alpha-band power A - FFT calculated EC-EO changes in alpha-band power (Δ*α*) vs fixed-order ARMA calculated changes B - FFT calculated EC-EO changes in alpha-band power vs a Lorentzian approximation based on the dominant alpha oscillatory mode calculated using the fixed order ARMA. For further details see Materials and Methods: Time series analysis in the main text. Results calculated for all EEG channels and participants.

**Figure S3:**
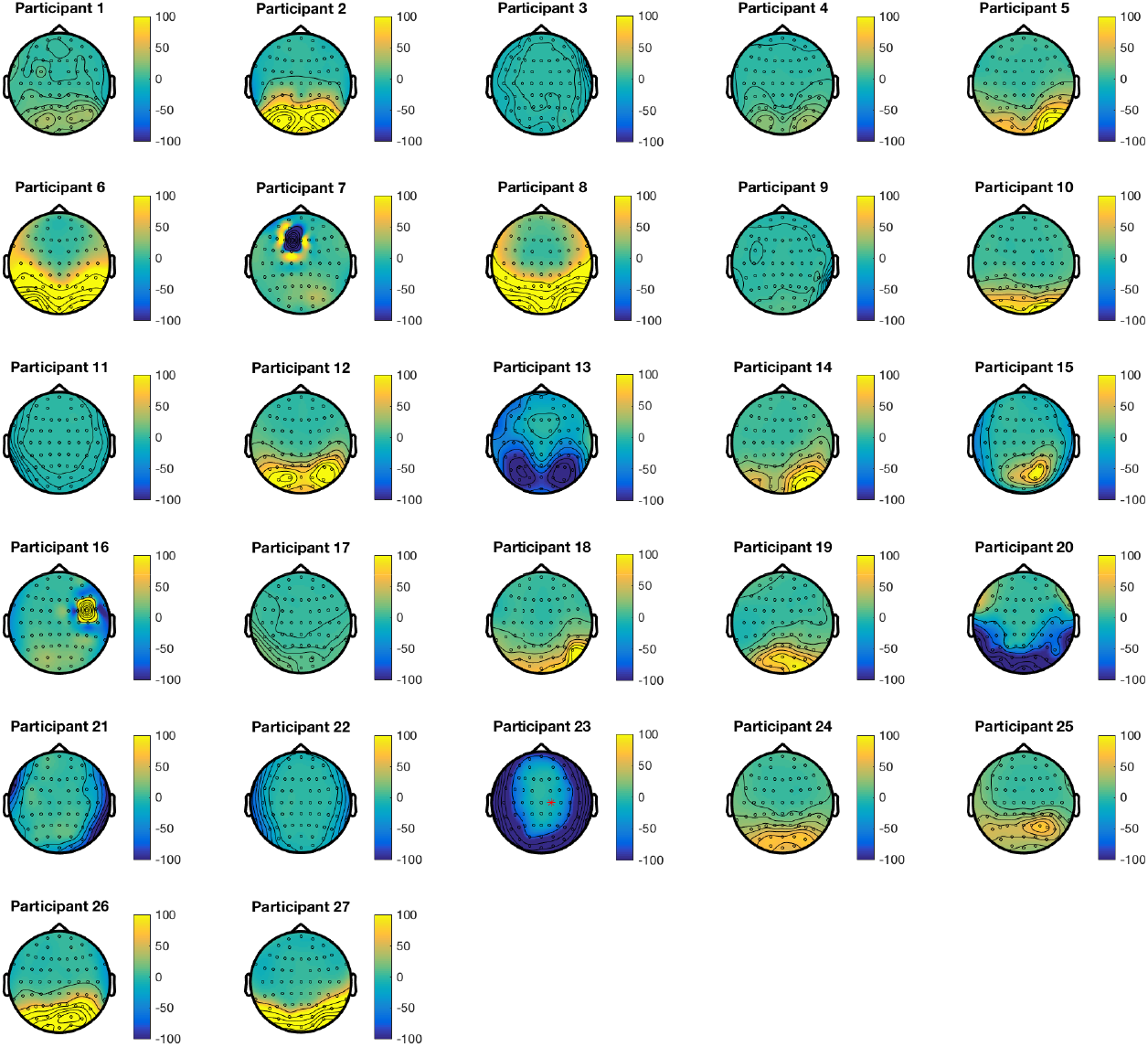
EC-EO changes in alpha (8-13 Hz) band spectral power for all 27 participants. The significant inter-participant variability should be noted. Scale bar units = *μV*^2^.

**Figure S4:**
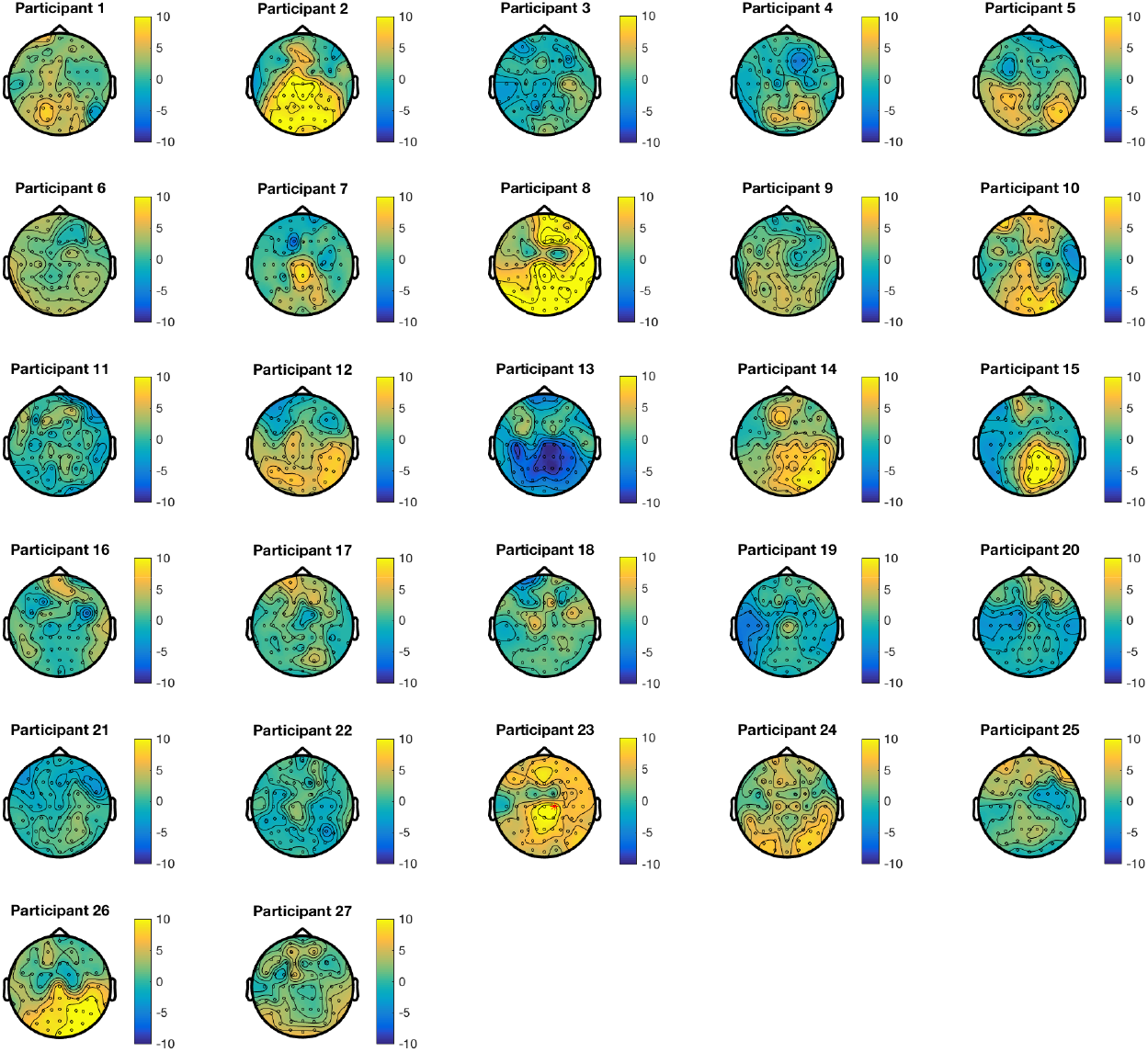
EO-EC changes in the damping of ARMA estimated alpha oscillatory mode activity for all 27 participants. Scale bar units = *s*^-1^.

**Figure S5:**
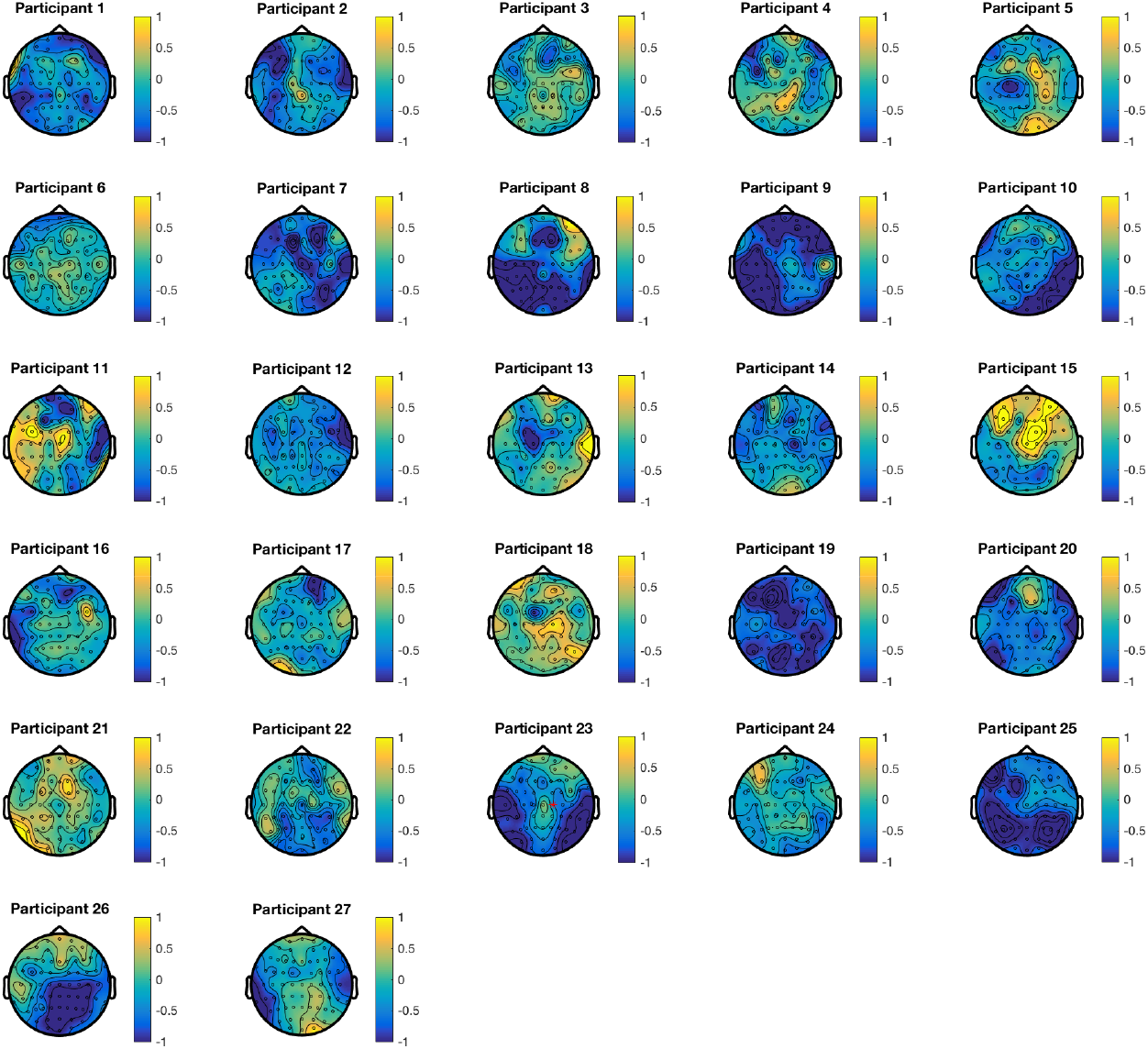
EC-EO changes in the frequency of ARMA estimated alpha oscillatory mode activity for all 27 participants. Scale bar units = Hz.

**Figure S6:**
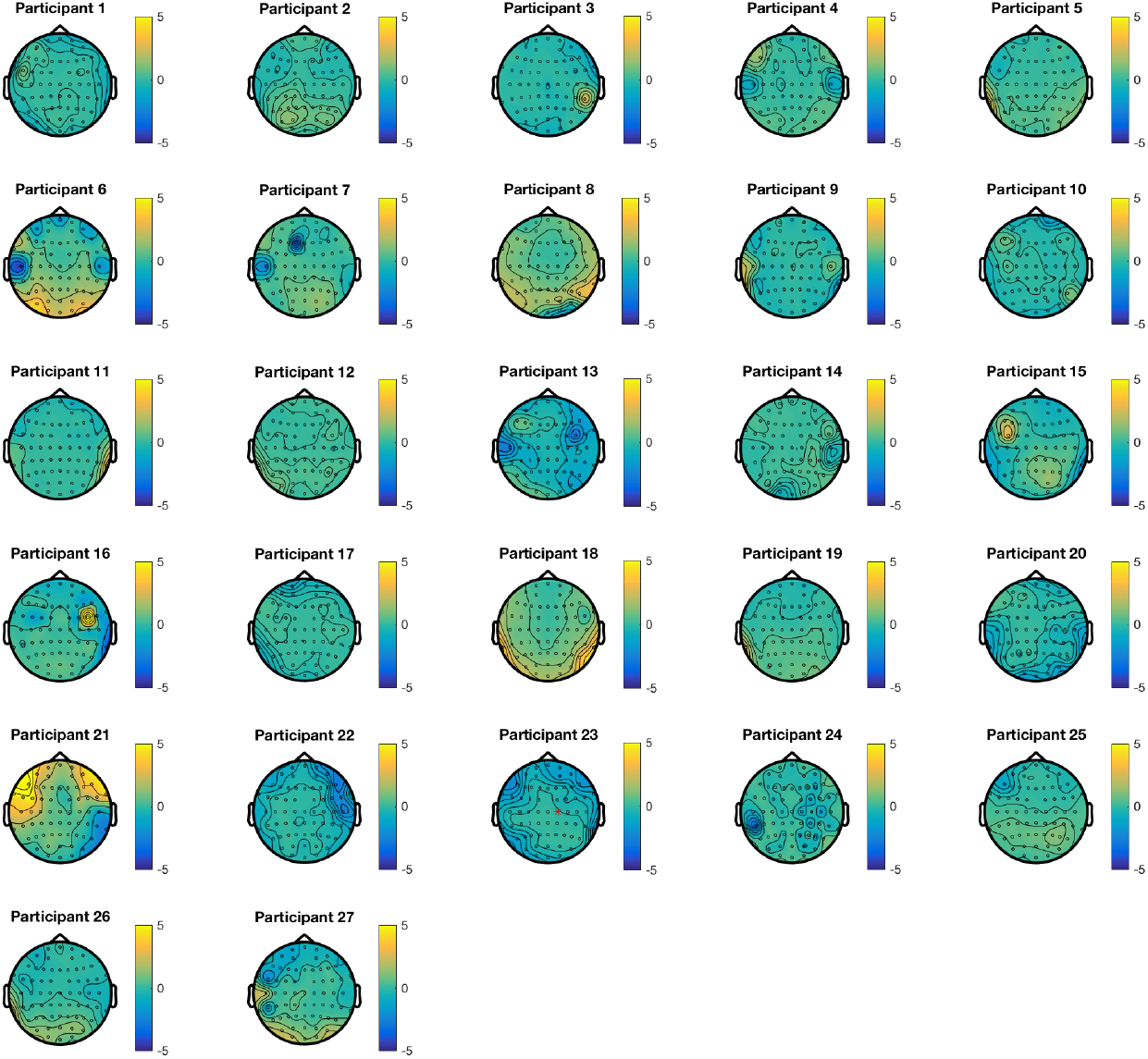
EC-EO changes in the estimated cortical input for all 27 participants. Scale bar units = *μV*.

**Figure S7:**
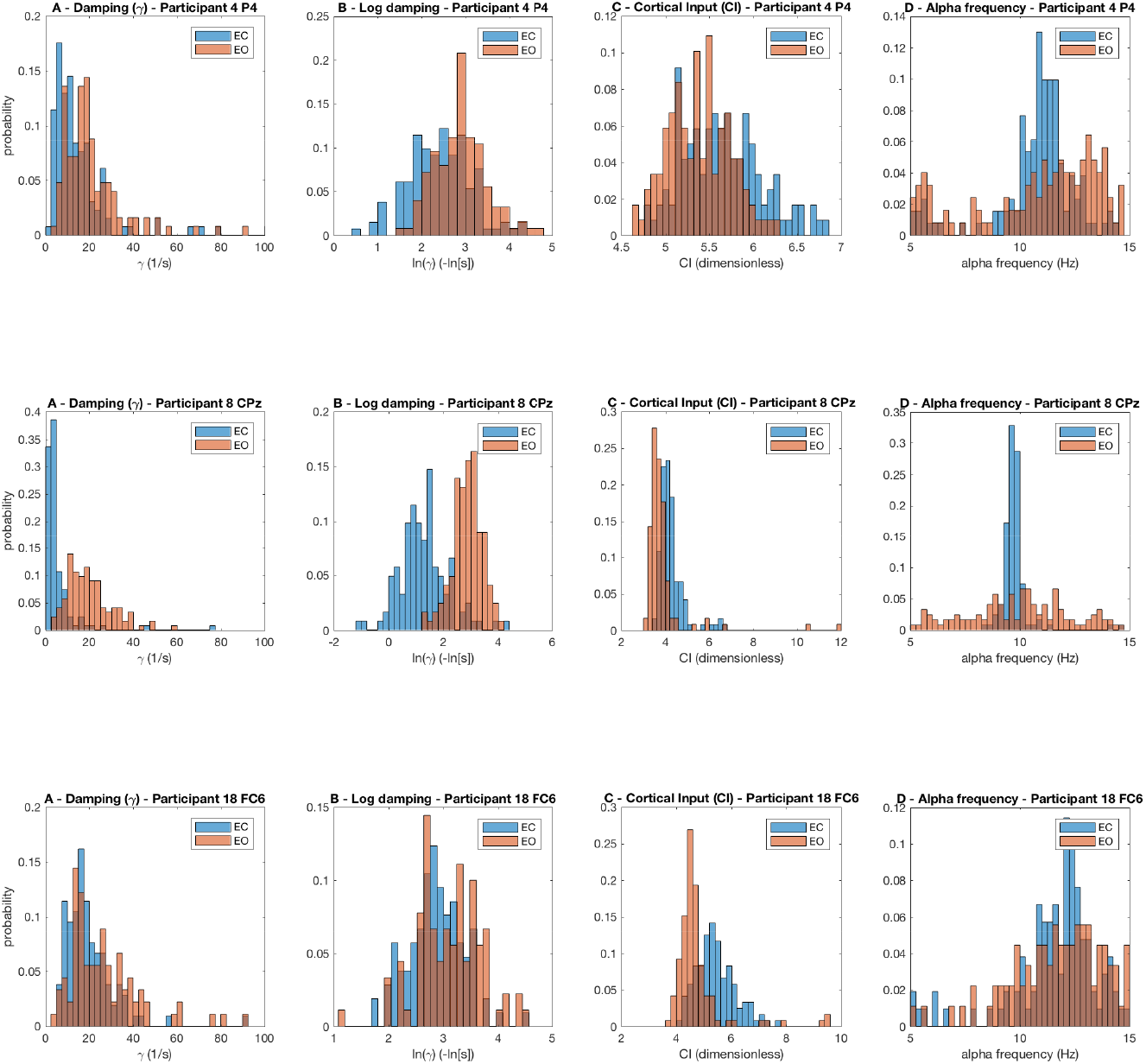
Representative distributions for the damping of ARMA estimated alpha oscillatory modes and cortical input for a number of individual subjects during EC and EO.

